# Necrotizing Soft Tissue Infections *Staphylococcus aureus* - but not *Streptococcus pyogenes*-isolates display high rate of internalization and cytotoxicity toward human myoblasts

**DOI:** 10.1101/530493

**Authors:** Jessica Baude, Sylvère Bastien, Yves Gillet, Pascal Leblanc, Andreas Itzek, Anne Tristan, Michèle Bes, Stephanie Duguez, Karen Moreau, Binh An Diep, Anna Norrby-Teglund, Thomas Henry, François Vandenesch, and INFECT Study Group

## Abstract

Necrotizing Soft Tissue Infections (NSTIs), often reaching the deep fascia and muscle, are mainly caused by group A *Streptococcus* (GAS) and to a lesser extent by *Staphylococcus aureus* (SA). Conversely SA is a leading etiologic agent of pyomyositis suggesting that SA could have a specific tropism for the muscle. To assess the pathogenicity of these two bacterial species for muscles cells in comparison to keratinocytes, adhesion and invasion of NSTI-GAS and NSTI-SA were assessed on these cells. Bloodstream infections (BSI) SA isolates and non-invasive coagulase negative Staphylococci (CNS) isolates were used as controls.

SA isolates from NSTI and from BSI exhibited stronger internalization into human keratinocytes and myoblasts than CNS or NSTI-GAS. While the median level of SA internalization culminated at 2% in human keratinocytes, it reached over 30% in human myoblasts due to a higher percentage of infected myoblasts (>11%) as compared to keratinocytes (<3%) assessed by transmission electron microscopy. Higher cytotoxicity for myoblasts of NSTI-SA as compared to BSI-SA, was attributed to higher levels of *psm*α and *RNAIII* transcripts in NSTI group as compared to hematogenous group. However, the two groups were not discriminated at the genomic level. The cellular basis of high internalization rate in myoblasts was attributed to higher expression of α5β1 integrin in myoblasts as compared to keratinocytes. Major contribution of FnbpAB-integrin α5β1 pathway to internalization was confirmed by isogenic mutants.

Our findings suggest the contribution of NSTI-SA severity by its unique propensity to invade and kill myoblasts, a property not shared by NSTI-GAS.

**Importance:** Necrotizing Soft Tissue Infection (NSTI) is a severe infection caused mainly by group A *Streptococcus* (GAS) and occasionally by *S. aureus* (SA); the latter being more often associated with pyomyositis. NSTIs frequently involve the deep fascia and may provoke muscle necrosis. The goal of this study was to determine the tropism and pathogenicity of these two bacterial species for muscle cells. The results revealed a high tropism of SA for myoblasts and myotubes followed by cytotoxicity as opposed to GAS that did not invade these cells. This study uncover a novel mechanism of SA contribution to NSTI with a direct muscle involvement, while in GAS NSTI this is likely indirect, for instance, secondary to vascular occlusion.

## Introduction

Necrotizing fasciitis and other necrotizing soft tissue infections (NSTIs) are the most extreme development of skin and soft tissue infections, requiring intensive care and surgical removal of necrotic tissues. NSTI encompass various clinical entities (1, 2). They are characterized anatomically by necrotic infections beyond the subcutaneous tissue and the superficial fascia, often involving the deep fascia and sometimes associated with myonecrosis (2). Based on the polymicrobial or monomicrobial character of the infection, NSTIs are classified as type I and type II, respectively. Type I is a polymicrobial infection involving aerobic and anaerobic organisms and, depending of the study, is either less or equally prevalent to type II (1). Type II is a monomicrobial infection caused by group A Streptococcus (GAS) and to a lesser extent by *Staphylococcus aureus* (SA), notably methicilin-resistant *S. aureus* (MRSA) (3). Although SA is a common cause of superficial SSTI and cellulitis (4), the exact prevalence of SA in NSTI is difficult to assess, principally because of the lack of prospective controlled population-based studies. Myositis has been reported as “an unexpected finding” in SA NSTI cases (3) and SA is typically an agent of pyomyositis, notably in tropical countries (5–8). Altogether, these epidemiological reports suggest a particular tropism and pathogenesis of SA for the skeletal muscles. The aim of the present study was to investigate and compare the invasiveness of muscle cells by SA and GAS as the two major species responsible for type II NSTI. We showed that SA displays a unique propensity to invade muscle cells principally via the FnbpAB-integrin α5β1 pathway, a property that is not seen in GAS from NSTI patients.

## Results

### SA display enhanced internalization rate in myoblasts as compared to other species

The capacity of both SA and GAS isolates to adhere and invade keratinocyte or myoblasts was evaluated using a gentamicin protection assay. NSTI-SA and bloodstream infections (BSI-SA, used as non-NSTI invasive infections controls) exhibited stronger adhesion than non-invasive coagulase negative Staphylococci (CNS) or NSTI-GAS, onto human keratinocytes (Fig 1A). Internalization into keratinocytes was also significantly higher for NSTI-SA, as compared to the other groups, although the frequencies of infected cells were relatively low (mean 2%) (Fig 1B). Adhesion on and internalization into murine and human myoblasts showed contrasting results. While there were no significant differences between NSTI-SA and NSTI-GAS in adhesion on murine and human myoblasts (Fig 2A and 2B), a dramatic difference in internalization rate was observed, reaching 30-40% for SA (NSTI and BSI) and only 1-5% for NSTI-GAS (Fig 2C and 2D). This very high level of internalization into murine and human myoblast is unique to SA and is shared by all SA strains of the present study. To determine if this high internalization level was caused by a higher mean number of infected cells or by a higher mean number of bacteria per cells, transmission electron microscopy (TEM) was performed on human myoblasts and keratinocytes upon infection by one NSTI-SA (INFECT2127) strain and one SA reference strain (SF8300). The results revealed that the mean number of bacteria per infected cells (actually TEM section) was equivalent in myoblast and keratinocytes (average 3-4 bacteria per infected cell) (Table 1).

**TABLE 1.** Quantification of intracellular bacteria assessed by transmission electron microscopy

| Cells | Strains | Number of analysed cells | Number of infected cells<sup>a</sup> | Number of bacteria in infected cells<sup>a</sup> | Mean number of bacteria per infected cells | Percentage of infected cells |
| --- | --- | --- | --- | --- | --- | --- |
| Myoblasts | INFECT2127 | 301 | 44 | 154 | 3,5 | 14.62 |
|  | SF8300 | 175 | 20 | 76 | 3.8 | 11.43 |
| Keratinocytes | INFECT2127 | 455 | 13 | 45 | 3.46 | 2.86 |
|  | SF8300 | 491 | 7 | 26 | 3.7 | 1.43 |

<sup>a</sup>Assessed by visual counting of TEM sections

**Fig. 1.**
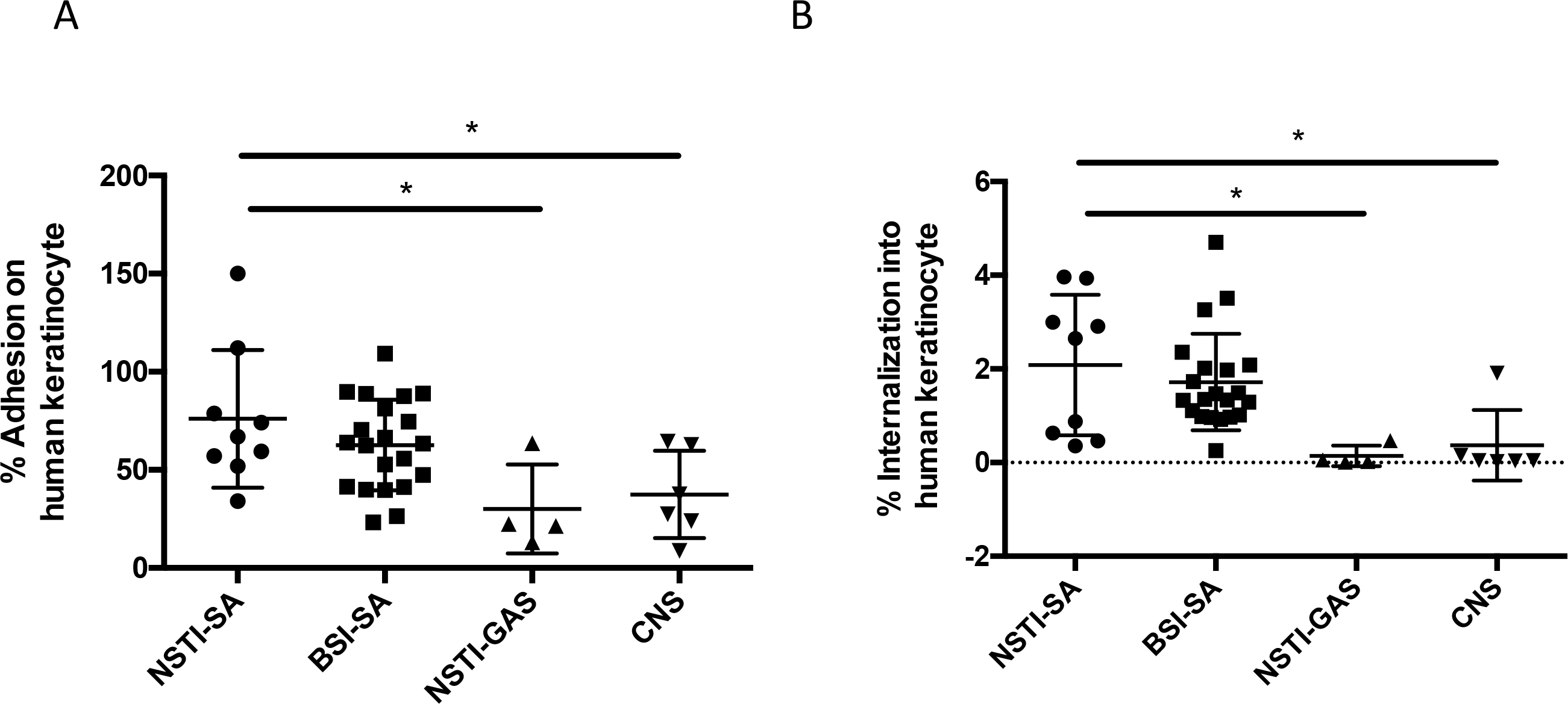
Adhesion and internalization of *Staphylococcus aureus* (SA), coagulase negative *Staphylococcus* (CNS) and group A *Streptococcus* (GAS) to human keratinocytes. (A) Percentages of adhered bacteria were calculated after 2 hours of infection in relation to inoculum of infection. Intracellular bacteria assessed in B were subtracted; (B) Internalization was determined after 2 hours of infection followed by antibiotic treatment to exclude extracellular bacteria. Multiplicity of infection (MOI) = 10. The horizontal lines within each group represent the mean value ± standard deviations of at least three independent experiments per strain. *p<0.05, **p<0.01, *** p<0.001.

**Fig. 2.**
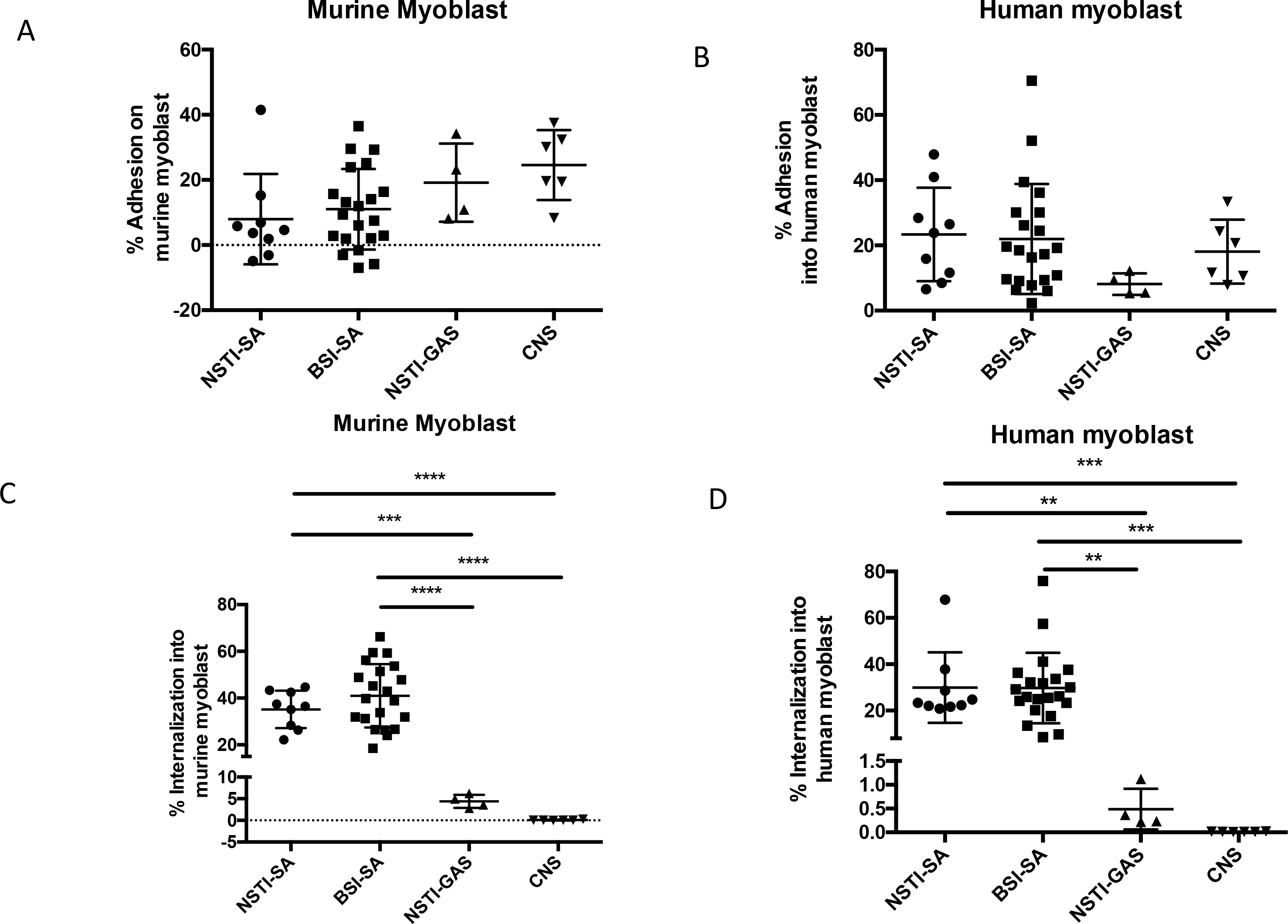
Adhesion and internalization of *Staphylococcus aureus* (SA), coagulase negative *Staphylococcus* (CNS) and group A *Streptococcus* (GAS) to murine and human myoblasts. (A and B) Percentages of adherent bacteria were calculated after 2 hours of infection in relation to inoculum of infection. Intracellular bacteria assessed in C and D were subtracted; (C and D) Internalization was determined after 2 hours of infection followed by antibiotic treatment to exclude extracellular bacteria. Multiplicity of infection (MOI) = 10. The horizontal lines within each group represent the mean value ± standard deviations of at least three independent experiments per strain. *p<0.05, **p<0.01, *** p<0.001, ****p<0,0001.

Conversely the percentage of infected cells was significantly higher in human myoblast (14.62 and 11.43% for INFECT2127 and SF8300, respectively) as compared to keratinocytes (2.86 and 1.43% for INFECT2127 and SF8300, respectively), which is in the order of magnitude of the difference in internalization rates between the two cell types as observed by gentamycin protection assay.

### The high rate of SA invasion into muscle cells is not an artifact of immaturity

To rule out the possibility that the observed phenotype in myoblasts could be an artifact of immature cells, the same experiments were performed in myoblast-derived myotubes. Maturity of the cells was assessed morphologically and also by the occurrence of contractibility under video microscopy (Movie S1-S2). In these conditions, the level of internalization of NSTI-SA and BSI-SA was 29.7% and 27.5% respectively, and thus not different to that obtained in myoblasts with the same strains (34.8% and 25.7% for NSTI-SA and BSI-SA respectively) (Fig. S1). These data indicate that the high internalization rate of SA is a shared property of the myoblastic lineage.

### FnbpAB is the major determinant of SA internalization into myoblast

To determine which *S. aureus* surface proteins are involved in SA internalization within myoblast, mutant strains for the genes encoding the main effectors of SA internalization, namely, FnbpAB, ClfA, Atl, SdrD, Tet38 and Gap (Table S1) (9), excluding those not present in all our SA strains (Eap), were tested. The results revealed a major contribution of FnbpAB. Indeed, while the single *fnb*A and B mutants had no significant phenotype, the double mutant delta-*fnb*AB was internalized 1000-fold less (internalization reduced by 99.999%) than the wild type (WT) strain (Fig. 3). The major impact of FnpbAB was confirmed by using another *fnb*AB double mutant in strain 8325-4 (DU5883) (not shown). Other adhesins also contributed to some extent to internalization into myoblast: mutants in *atl, gap, clf*A and *tet*38 were impaired in internalization by 71%, 27.4%, 25% and 24.6 %, respectively, as compared to the WT strain. (Fig. 3).

**Fig. 3.**
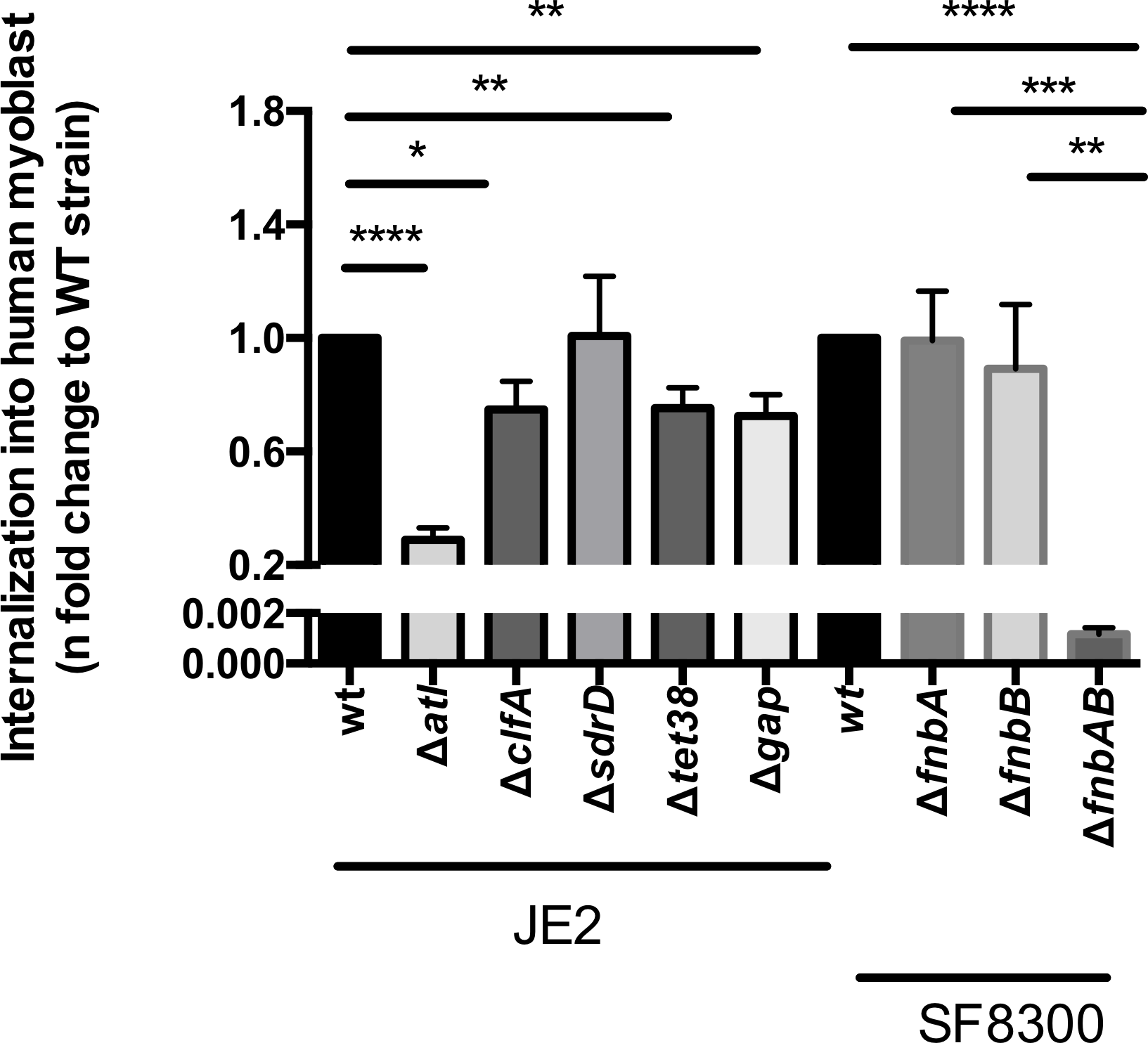
Internalization of *S. aureus* mutants into human myoblasts. Internalization of the various mutants in genes encoding surface proteins was determined after 2 hours of infection followed by antibiotic treatment to exclude extracellular bacteria. The internalization was normalized to that of wild type strain. Multiplicity of infection (MOI) = 10. All values are means ± standard error of mean of three independents experiments in duplicate for each strain. *p<0.05, **p<0.01, *** p<0.001, ****p<0,0001.

### Myoblasts express increased level of Integrin α5β1

To investigate the cellular basis of high internalization rate of SA in myoblasts (ca 30-40%) as compared to keratinocytes (ca 2%), we analyzed the expression of integrin α5β1 (the major target of FnbpAB internalization pathway) on human keratinocyte and myoblasts. Flow cytometry analysis by using an anti-α5β1 antibody showed a greater expression of the α5β1 integrin at the surface of myoblasts as compared to keratinocytes (Fig. 4A). Next, the transcription of integrin α5β1 as well as other eukaryotic surface receptors involved in *S. aureus* internalization (9) was evaluated by RT-qPCR in the two cell lines. The integrin α5β1 displayed contrasting results: whilst integrin β1 was expressed at the same level in both cell lines, integrin α5 was expressed three times more in myoblasts. The expression of integrin β1 was at least 20 times higher than integrin α5, implicating integrin α5 as the limiting partner of the heterodimer (Fig. 4B). Altogether, flow cytometry and RT-qPCR results suggest that functional α5β1 is expressed at higher level in myoblasts as compared to keratinocytes. Finally, the major role of integrin α5β1 pathway for SA internalization into myoblasts was confirmed by preincubation of cells with anti-α5β1 antibodies before being challenged by SA, which reduced internalization by more than 65% (Fig. S2). In addition, the transcription of all other cellular receptors known to be involved in SA internalization (IntegrinαV, Integrinβ3, Hsc70, Annexin 2, Hsp60, CD36, gp340) (9) was studied. Increased expression tendancy was generally observed in myoblast as compared to keratinocytes (Fig. S3). Altogether, these results strongly suggest that the higher internalization of SA in myoblast compared to keratinocytes is mediated by various pathways with a preeminence of the FnBP-Fibronectin-α5β1 integrin-mediated pathway.

**Fig. 4.**
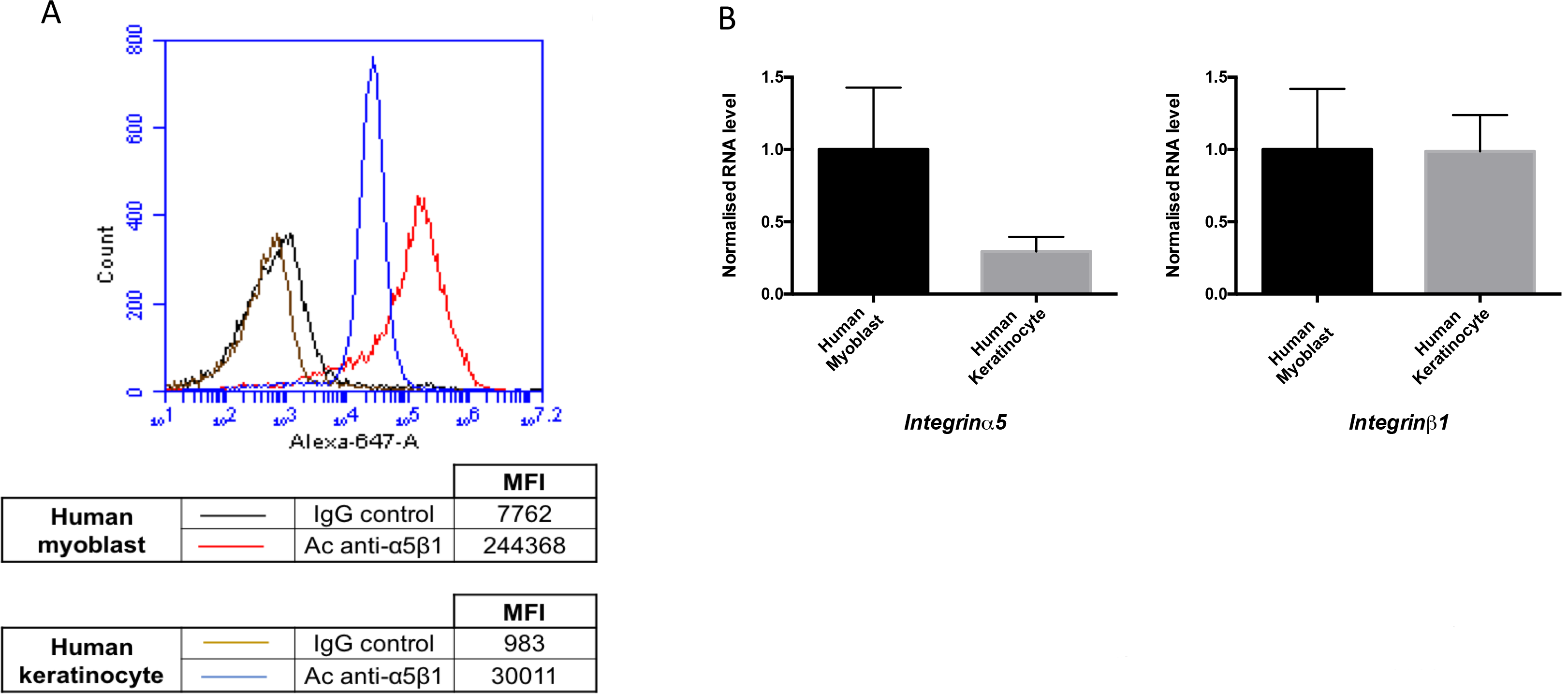
Expression of α5β1 integrin by human keratinocyte and myoblasts. (A) Cell surface expression of α5β1 integrin assessed by flow cytometry and α5β1 antibody; (B) Relative transcript levels of α5, β1 integrin sub-units were determined using quantitative reverse-transcriptase PCR, normalized to the internal *β-actin* standard. All values are means ± standard error of mean of four independents experiments.

### Intracellular *RNAIII* and *psm*α transcript levels correlate with cytotoxicity of NSTI-SA

Since the level of internalization in a given cell type does not inform about the fate of internalized bacteria (i.e. dormancy versus cytotoxicity), the toxicity associated with bacterial internalization was estimated at 24h post-infection by the lactate dehydrogenase (LDH) quantification. NSTI-SA and BSI-SA were as cytotoxic as GAS on keratinocytes; of note CNS were significantly less cytotoxic toward this cell line than other bacterial groups. The most noticeable cytotoxic effect of NSTI-SA was observed on murine and human myoblasts as compared to all other groups, namely NSTI-GAS, CNS, and unexpectedly BSI-SA. (Fig. 5). Of note, the culture supernatant from SA and GAS cultures did not produce cytotoxicity on keratinocytes and myoblasts (Fig S4), assessing that the observed cytotoxicity was not caused by toxins or other products released by the bacteria. To elucidate the basis of higher cytotoxicity for myoblasts of NSTI-SA as compared to BSI-SA, the level of the major effectors of intracellular *S. aureus* cytotoxicity, namely PSMα, Hla, Ldh and Agr-RNAIII (10, 11) were assessed by RT-qPCR. The levels of *RNAIII* and *psm*α were significantly higher in NSTI-SA strains as compared to BSI-SA in intracellular conditions into human myoblasts (Fig. 6) as well as from planktonic culture (data not shown). In contrast, the level of *hla* and *ldh* transcript were not significantly different between the two groups.

**Fig. 5.**
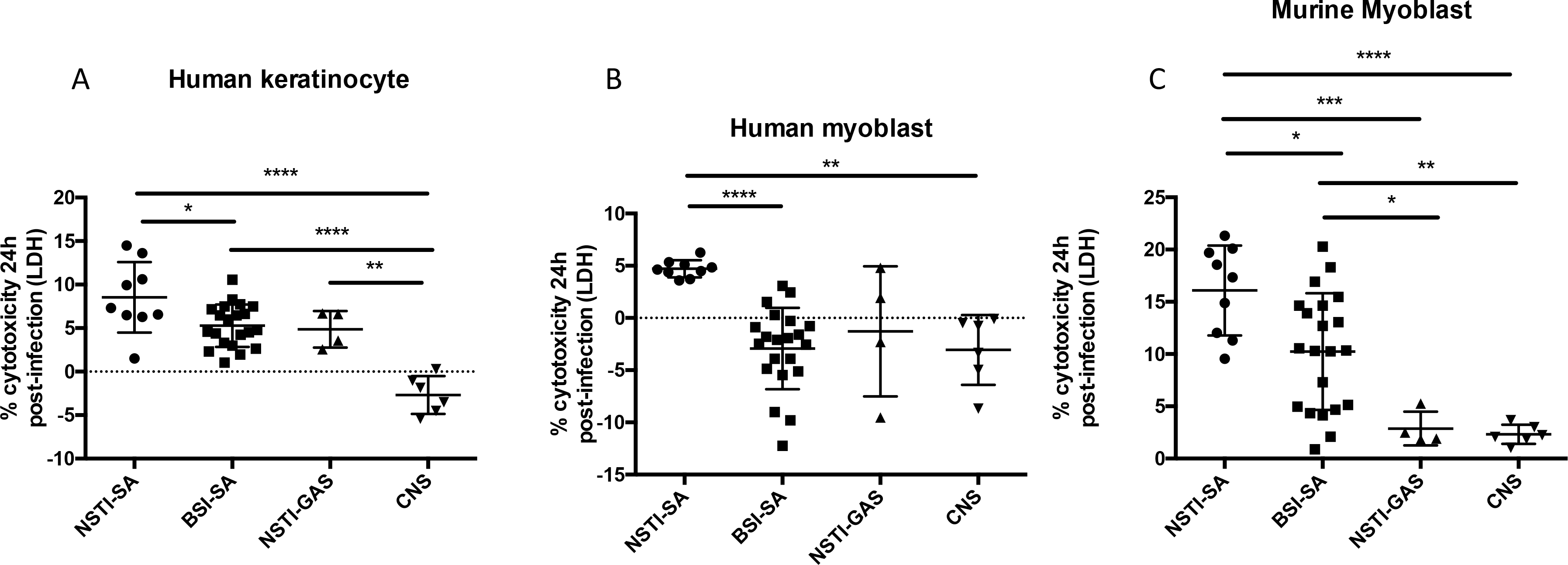
Cytotoxicity of intracellular bacteria at 24h post infection. Cytotoxicity was estimated by quantifying LDH release by infected cells at 24 hours post infection. Multiplicity of infection (MOI) = 10. The percentage of cytotoxicity was calculated as follows: ((LDH infected cells − LDH lower control)/(LDH higher control − LDH lower control)) ×100. The horizontal lines within each group represent the mean value ± standard deviations of at least three independent experiments per strain. *p<0.05, **p<0.01, *** p<0.001, ****p<0,0001.

**Fig. 6.**
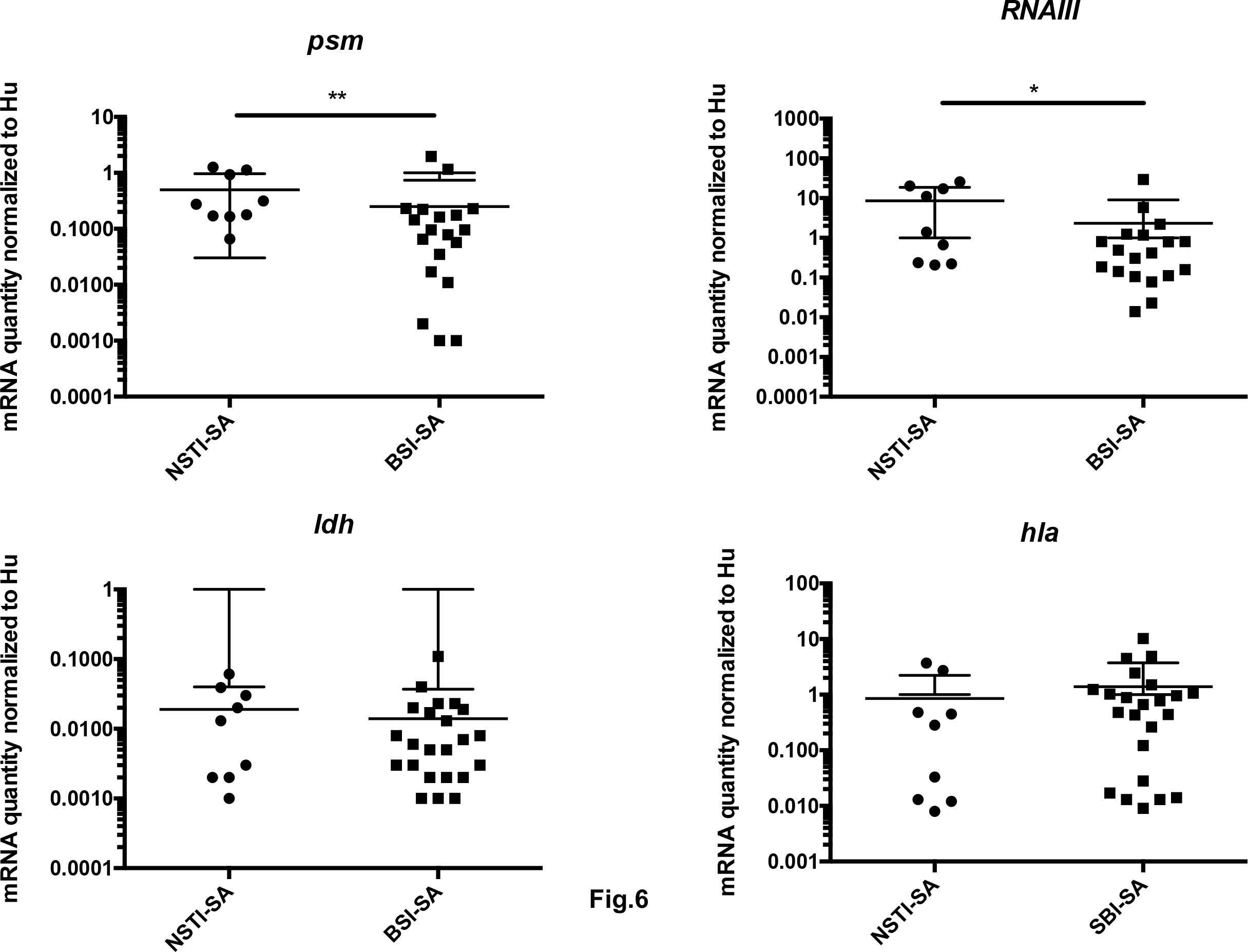
Expression of intracellular bacterial mRNA into human myoblasts 3 hours post infection. Relative transcript levels were determined using quantitative reverse-transcriptase PCR and expressed as n-fold change to the internal *hu* standard. The horizontal lines within each group represent the mean value ± standard deviations. *p<0.05, **p<0.01.

### Differences in cytoxicity between NSTI-SA and BSI-SA is not resolved at the genomic level

In order to confirm that the differences between NSTI-SA and BSI-SA isolates in *RNAIII* and *psm*α expression were not related to a genomic bias, a genomic analysis was conducted on the 30 *S. aureus* strains. Phylogenetic analysis based on core-genome confirmed that NSTI and hematogenous strains were phylogenetically entangled (Fig. S5). Virulence factor distribution assessed by micro-array or WGS was not contributive to distinguish NSTI-SA from BSI-SA isolates (Table 2). Notably, there was no virulence factors discriminating the two groups. Detailed genomic could not identify discriminant marker between the two groups either considering genomic determinants presence/absence (including ncRNA, virulence genes,…) or single nucleotide polymorphism (SNPs) (Text. S1).

**TABLE 2.**
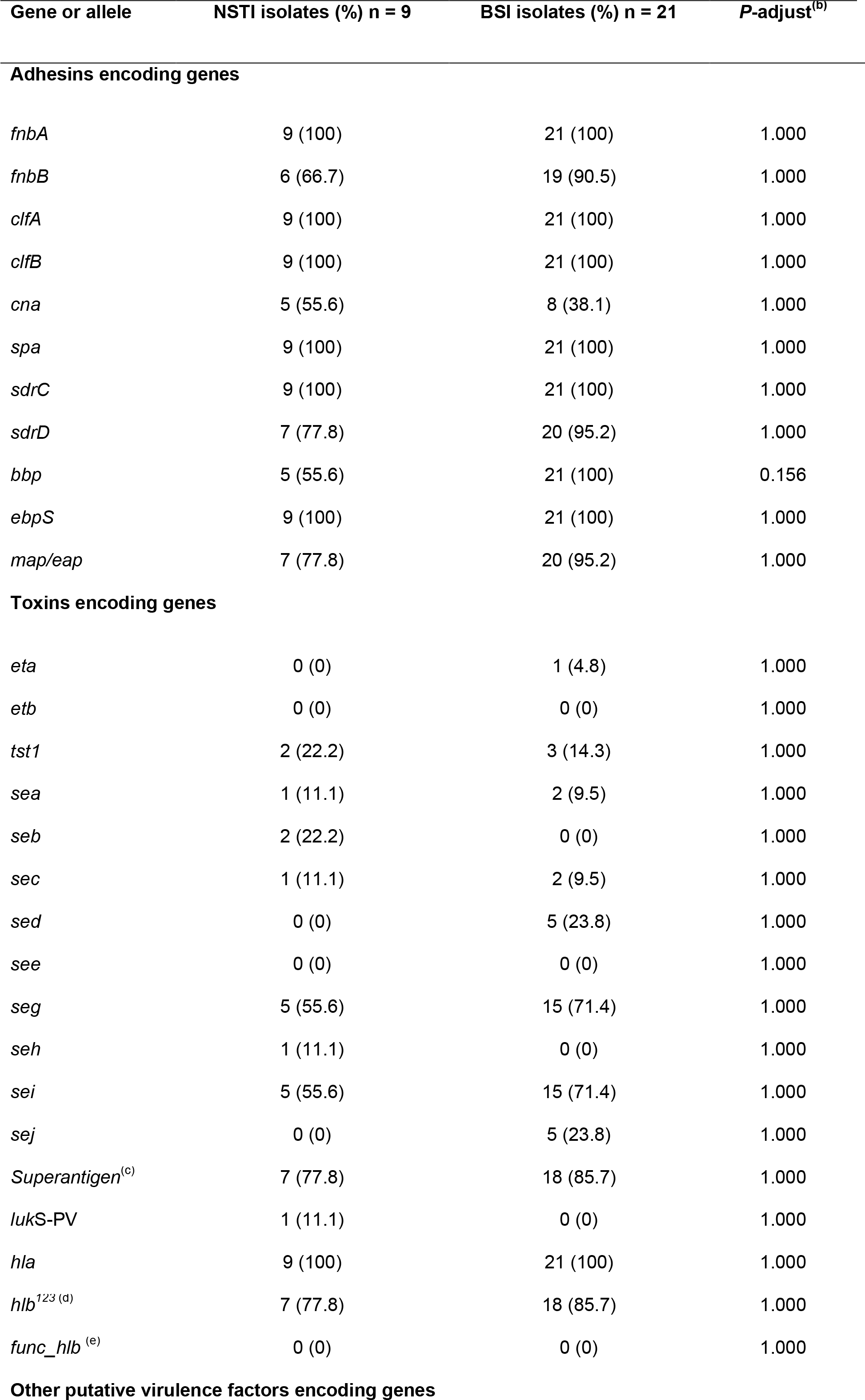
Frequency of the genes detected by DNA microarray in *S. aureus* NSTI and BSI isolates^a^.

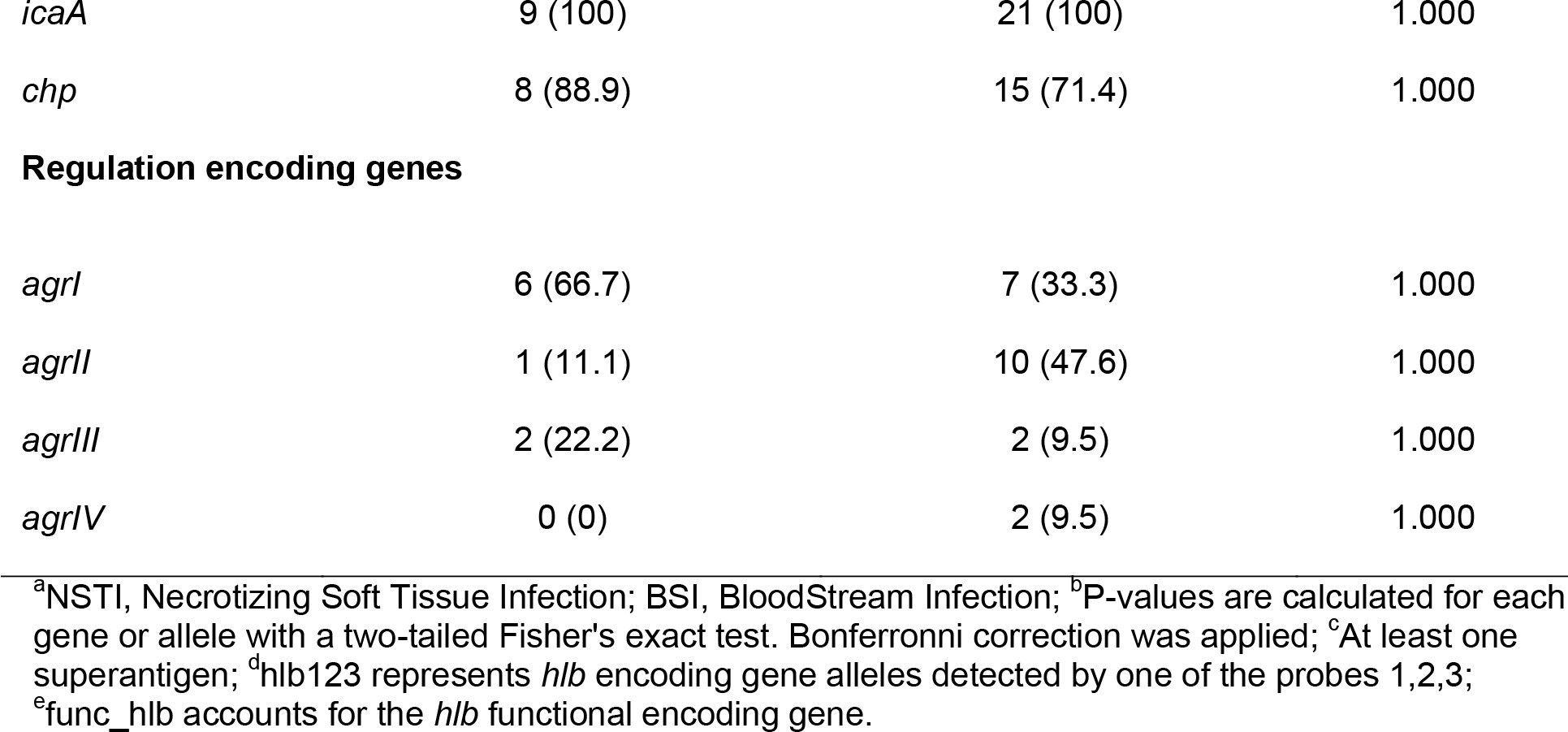

## Discussion

Necrotizing fasciitis is a rare disease that involves superficial fascia and results in the extensive damage and necrosis of the surrounding tissue including muscle. In the present study we show that SA may contribute to this disease by its unique propensity to invade muscle cells and consequently trigger cell death, a property that is not seen in GAS from NSTI patients. Interestingly, this unique capacity of *S. aureus* to invade myoblast and myotube from both mice and human origin is uncommon, as neither CNS nor GAS did internalize in those cells to such efficiency. Moreover, to our knowledge, there is no other type of eukaryotic cells in which SA internalizes at this level (over 30%), e.g. the level of SA internalization is ca. 17% in human embryonic kidney cells (12), 8% in endothelial cells (12), 2% in human alveolar epithelial cells (A549) (K. Moreau, unpublished data), <1% HELA epithelial cells (13), <1% within osteoblast (14) and <2% in osteoclast (15). We could demonstrate that this high rate of internalization depends on several of the known pathways of *S. aureus* internalization with a preeminence of the FnBP-Fibronectin-α5β1 integrin-mediated pathway. The increased expression at the surface of human myoblasts of α5β1 integrin and other known receptors involved in SA internalization (integrin αV and β3, HSC70, Annexin 2, Hsp60, CD36 and Gp340) is a likely explanation for the high internalization rate of SA within myoblasts as compared to keratinocytes. This model is also consistent with TEM observation of a higher percentage of infected cells as compared to keratinocytes. The direct consequence of this high rate of internalization within myoblast is a higher cytotoxicity to these cells, and not, as observed in other setting a reduced cytotoxicity associated with high internalization rate (11).

Virulence factors that could be associated with NSTI-SA have been largely debated, Panton Valentine Leucocidin being frequently reported in US studies involving community-acquired MRSA USA300 (3). Other cases associated with *tst*- (16) or with *egc*-encoding isolates were reported (17), reminiscent of the involvement of superantigens in severe GAS tissue infections (18). Noticeably, the 9 NSTI-SA strains of the present study were not clonally related; none of these were MRSA and a single one was PVL-positive (Table 2). Their genetic content regarding virulence factors including superantigens was not significantly different from the BSI-SA strains that represent another major invasive disease (Table 2). WGS comparison by multiple approaches could not discriminate the two diseases isolate groups (Text S1). This lack of discrimination might be caused by the small size of the NSTI group leading to insufficient statistical power. However, the observation that BSI-SA were less cytotoxic than NSTI-SA with similar internalization rate (Fig. 5), is in accordance with previous studies showing that skin and soft tissue infection isolates produce more PSMα (as observed in our study with NSTI isolates) than infective endocarditis or hospital-acquired pneumonia isolates (19). This is also reminiscent of previous observation suggesting an association between low-toxicity and bacteraemia (20): It has been hypothesized that such invasive bloodstream infections limit the opportunities for onward transmission, whilst highly toxic strains associated with skin infections could gain an additional between-host fitness advantage, potentially contributing to the maintenance of toxicity at the population level (20). Altogether this analysis reveal that despite similar virulence factor repertoire, NSTI-SA and BSI-SA isolates differ in their phenotypic expression of virulence and may thus be considered as different pathovars.

The respective contribution of GAS and SA to NSTI is a matter of debate. Of note, the four strains of GAS isolated from NSTI patients, despite being isolated from very severe cases of NSTI, did not present a strong invasive capacity for muscle cells *in vitro*. This result is counter-intuitive given the involvement of epidermis, dermis, subcutaneous tissue, fascia, and muscle in GAS-NSTI (1). This suggests that the muscle involvement in GAS NSTI does not result from bacterial invasion of muscle cells whilst it could be the cases for SA-NSTI when muscle involvement occurs as described (3). However, if the muscle destruction seen in GAS-NSTI is not caused by direct intracellular toxicity of GAS, it is likely not caused either by the toxicity of secreted enzymes and toxins for the muscle cells, as culture supernatant neither from GAS nor from SA had any significant toxicity on cultured myoblasts. It is thus more likely that muscle necrosis in GAS-NSTI results from vascular occlusion secondary to platelet-leukocytes aggregation in response to infection of the adjacent anatomical skin and soft tissue structures.

In conclusion, our results uncover new insight into the pathophysiology of GAS and SA-NSTI and provide a new cellular basis for the contribution of *S. aureus* to pathogenesis of NSTI and other deep-seated infection involving the muscle. It also sustains the potential synergy of GAS and SA in NSTI, a situation that is not uncommon (2, 21). It advocates for future prospective clinical study systematically depicting the anatomical tissue layers of infection according to the micro-organisms.

## Material and Methods

### Ethics statement

Muscle biopsies for myoblast cell line were obtained from the Bank of Tissues of Research (Myobank BTR, a partner in the EU network EuroBioBank) in accordance with European recommendations and French legislation. Muscle biopsies for experiment in primary myoblast were obtained from NeuroBioTec (Hospices Civils de Lyon, *Centre de Ressources Biologiques*) and declared to the Ministry of Research under the agreement DC 2011-1437 and the number of accession AC 2012-1867.

### Bacterial strains

Bacterial isolates are listed in Table 3. They include strains of *S. aureus* (n=4) and *S. pyogenes* (n=4) isolated from patients included in the INFECT (Improving Outcome of Necrotizing Fasciitis: Elucidation of Complex Host and Pathogen Signatures that Dictate Severity of Tissue Infection) cohort which is an international, prospective, multicenter observational research project (Clinical Trials #NCT01790698). In addition, 5 *S. aureus* strains from French NSTI patients spontaneously referred to the French National Reference Laboratory for Staphylococci were included. The diagnosis of NSTI was made by the surgeon during the primary operation. It is based on, but not restricted to, findings of necrosis, dissolved or deliquescent soft tissue, and fascial and muscular affection (22). The invasive non-NSTI SA control strains consisted in Bloodstream infection (BSI) strains (12 infective endocarditis, 9 bacteremia without infective endocarditis) collected during a prospective cohort study on *S. aureus* bacteremia (VIRSTA) (23). Those strains were chosen in order to genotypically match NSTI strains on the basis of micro-array-deduced clonal complexes (23) confirmed by whole genome sequencing (Illumina MiSeq technology, Illumina, San Diego, USA, with 16-266 x coverage, median 137). Non *aureus*-staphylococci were type strains of the most frequently isolated CNS species. Strains USA300 JE2 and several transposons insertion mutants in genes encoding proteins potentially involved in internalization (*atl*, *clfA*, *sdrD*, *tet38*, *gap*, *strA*, *strB*) were retrieved from the public Nebraska library (24) with substantial help of Ken Bayles. In-frame deletion of the *fnb*A, *fnb*B and double mutant in the SF8300 background were performed as described previously (25, 26) using the pKOR1 allelic replacement mutagenesis system and the primers shown in Table S2 in the supplemental material. Laboratory strains and plasmids are listed in Table S1. All strains were cultured in Brain-Heart infusion (BHi Becton Dickinson) broth under agitation (200 rpm) at 37°C.

**TABLE 3.**
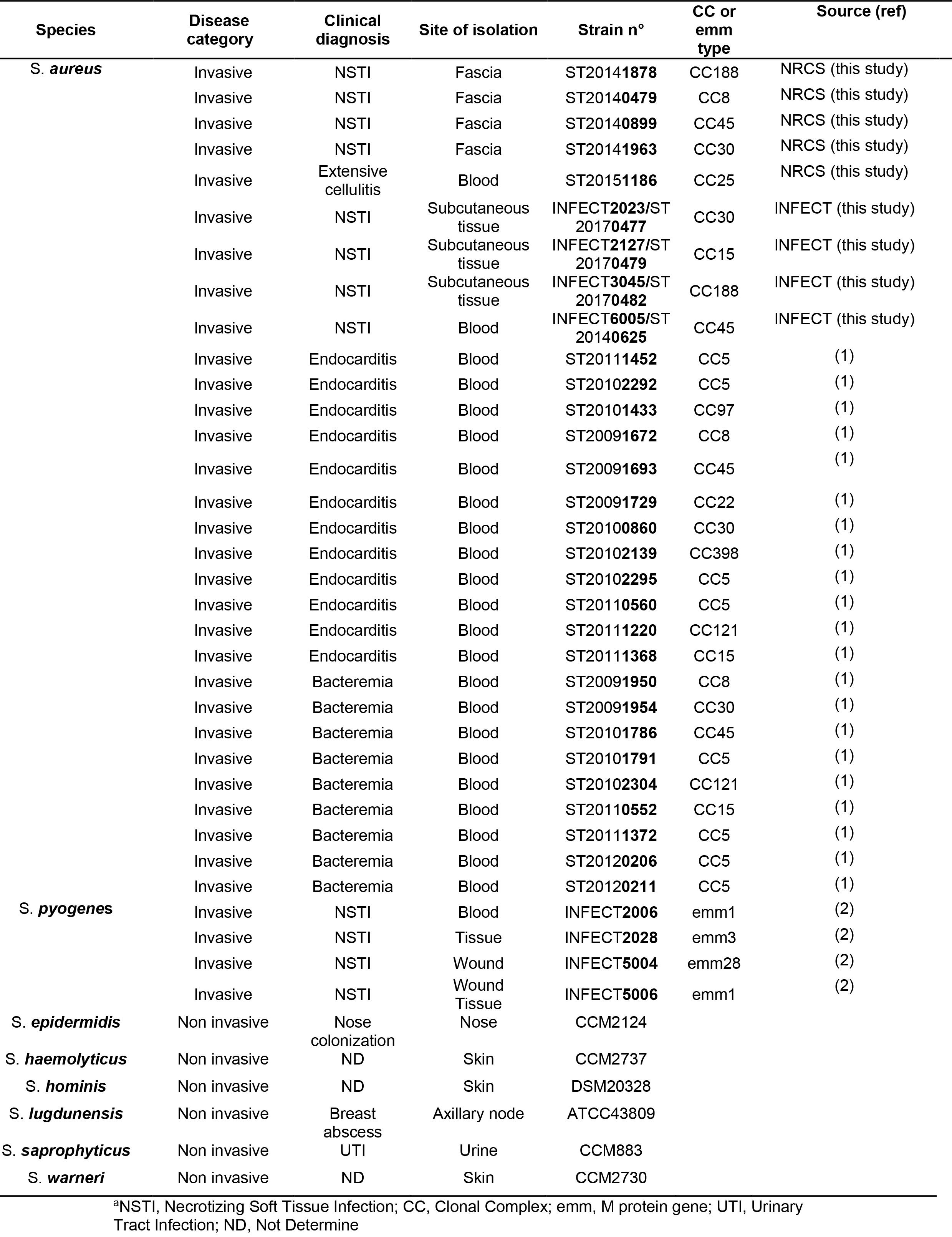
Description of clinical strains used in this study^a^

### Cells and culture conditions

The human keratinocytes N/TERT-1 cells (27) were cultured in keratinocyte SFM media (Thermofisher) at 37°C under 5% CO_2_ atmosphere. The murine myoblast cells C2C12 (28) were cultured in Dulbecco’s Modified Eagle’s Medium (Thermofisher) supplemented with 20% Fetal Bovine Serum (Gibco) and 1% penicillin/streptomycin. Murine myoblasts were differentiated into myotubes by addition of 2% of horse serum (Sigma) in the medium. The human myoblast line was obtained as follows. Deltoid muscle biopsy from a 71-year-old male was obtained from the Myobank BTR (see ethical above). Muscle stem cells were extracted, sorted and tested for their myogenicity as previously described (29). Cells were then expended in proliferating KMEM medium (1 volume of M199, 4 volumes of Dulbecco’s modified Eagle’s medium (DMEM), 20% fetal bovine serum (v/v), 25 μg/ml Fetuin, 0.5 ng/ml bFGF, 5 ng/ml EGF, 5 μg/ml Insulin). The lentiviral vector particles were produced by transient transfection of the packaging construct (HIV-1 psPAX2), a minimal genome (HIV-1 pLv-hTERT-CDK4 from CloneSpace LLC ref SKU: CS1031) bearing the expression cassettes encoding the Htert and CDK4 proteins and the puromycin resistance, and the VSV-G-envelope expressing plasmid pMDG2 (DNA ratio 8:8:4 μg) into 293T cells (3.5 × 10^6^ cells plated 1 day before transfection in 100-mm dishes) by the calcium phosphate method (30). Viral particles were normalized by an exogenous reverse transcriptase assay and titrated on human primary myoblast cells (31). Transductions of human primary myoblasts were carried out with different MOI in 12 well plates (80,000 cells plated 1 day before) in presence of 6 μg/ml of polybrene overnight in KMEM culture medium as previously described by Thorley (32). One day after transduction, cells were passaged and cultured in presence of puromycin (1 mg/ml) during 3-4 days until the death of non-transduced primary myoblast control cells was completed. Once puromycin selected, transduced myoblasts were cultured in KMEM-D medium (KMEM medium with dexamethasone 0.2 mg/ml) in absence of puromycin. After one week of culture in KMEM-D medium, immortalized myoblasts could be cultured for long term.

### Adhesion, internalization, survival and cytotoxic assays

The intracellular infection of cells was performed using gentamycin protection assay as described elsewhere with modifications (33). Cells were seeded at 80,000 cells/well in 24-well plates and incubated at 37°C with 5% CO_2_ for 24 h in culture medium. Bacterial cultures (9h of growth) were washed with PBS and resuspended in antibiotic-free culture medium at a concentration corresponding to a MOI of 10. The MOIs were subsequently confirmed by CFU counting upon agar plate inoculation. Cells were washed twice in PBS to remove antibiotics, and normalized bacterial suspensions were added to the wells. The infected cultures were incubated for 2h at 37°C and then washed with PBS. The cells used to assay invasion were incubated for an additional 1h at 37°C in culture medium containing 200 mg/L gentamicin and 10 mg/L lysostaphin to rapidly kill extracellular but not intracellular bacteria. To assay bacterial persistence, the cultures were further incubated in medium containing 40 mg/L gentamicin and 10 mg/L lysostaphin for 24h. These lower concentrations resulted in the killing of bacteria cells released upon host cell lysis, thus preventing infection of new host cells. Cells used to evaluate adhesion (2h post-infection), invasion (3h post-infection) and persistence (24h post-infection) were lysed by osmotic shock in sterile pure water, and extensively pipetted to achieve the full release of bacteria. The bacterial suspension was plated on Trypcase Soy Agar (TSA; bioMérieux, Marcy l’Etoile, France). After 24h of incubation at 37°C, the colonies were enumerated using an EasyCount automated plate reader (AES Chemunex). Infected cell death was quantified by the release of the cytosolic enzyme LDH into the culture supernatant at 24h post-infection using the CytoTox-ONE Homogeneous Membrane Integrity Assay (Promega) according to the manufacturer’s instructions. LDH release into the supernatant of infected cells was compared to that of uninfected cells that were either left intact (lower control) or fully lysed by osmotic shock (higher control). The percentage of cytotoxicity was calculated as follows: ((LDH infected cells − LDH lower control)/(LDH higher control − LDH lower control)) ×100.

### Relative quantification of bacterial RNA by RT-qPCR

Cells lines were infected as described above. After the first antibiotic treatment, cells and bacteria were harvested by trypsin detachment and centrifugation. Pellet was treated with 20 μg lysostaphin (1 mg/ml) and RNA isolation was performed using the RNeasy Plus mini kit (QIAGEN) according to the manufacturer’s instructions. The RNA was quantified using a NanoDrop spectrophotometer, and 100 ng of total RNA was reverse transcribed into cDNA using Reverse Transcriptase System (Promega). Two μL of 1/5 diluted cDNA was used as a template for the real-time PCR amplification using PowerUp SYBR^®^ Green Master Mix and a StepOne Plus system (Applied Biosystem) with specific primers shown in Table S2. Genes expression analysis was performed by using ΔCt methods using *hu* gene as an internal standard and confirmed by *gyr*B gene.

### Relative quantification of eukaryotic receptors by RT-qPCR

Total RNA was isolated from CTi400 and N/TERT-1 cells using RNeasy Plus mini kit (QIAGEN) according to the manufacturer’s instructions. Complementary DNA (cDNA) was synthesized from 1 μg of total RNA using random primers and Reverse Transcriptase System (Promega). The cDNA was used as a template for the real-time PCR amplification using PowerUp SYBR^®^ Green Master Mix and a StepOne Plus system (Applied Biosystem) with specific primers shown in Table S2. Genes expression analysis was performed by using ΔCt methods using β-*actin* gene as an internal standard and confirmed by GAPDH.

### Transmission Electron Microscopy

Infected cells were fixed in glutaraldehyde 2% and washed three times in saccharose 0.4M and sodium cacodylate buffer 0.2M pH 7.4 for 1h at 4°C, and postfixed with 2% OsO4 and Na C-HCl Cacodylate 0.3M pH 7.4 30 minutes at room temperature. Cells were then dehydrated with an increasing ethanol gradient (5 minutes in 30%, 50%, 70%, 95%, and 3 times 10 minutes in absolute ethanol). Impregnation was performed with Epon A (50%) plus Epon B (50%) plus DMP30 (1.7%). Inclusion was obtained by polymerisation at 60°C for 72h. Ultrathin sections (approximately 70 nm thick) were cut on a ultracut UC7 (Leica) ultramicrotome, mounted on 200 mesh copper grids coated with 1:1,000 polylisine, stabilized for 1 day at room temperature and, contrasted with uranyl acetate. Sections were examinated with a Jeol 1400JEM (Tokyo,Japan) transmission electron microscope equipped with a Orius 600 camera and Digital Micrograph. Cells were visually examined (175 myoblasts, 455 keratinocytes) to assess the number of infected cells and the number of bacteria per infected cells.

### Flow cytometry

Cells were trypsinized, washed and suspended in PBS at a concentration of 1.10^6^ per ml, and aliquots were incubated with Fc Block (ThermoFisher Scientific 14-9161-71) 20 minutes at 4°C. The primary α5β1 integrin antibody (10 μg/mL) (Merck Millipore MAB1999) was added for 30 minutes at 4°C. Cells were washed, resuspended in PBS before incubation with Alexa Fluor 647 secondary antibody (10 μg for 10^6^ cells) (Themofisher Scienfitic A-21236) for 30 minutes at 4°C and then washed 2 times in PBS. Finally, cells were assessed using a BD Accuri-C6 (BD Biosciences, Le pont de Claix, France) flow cytometer.

### Statistical analysis

The statistical analyses were performed using GraphPad Prism 6 software. Data from two groups were compared using either Student’s 2-tailed t-test for paired samples or Mann Whitney test for unpaired samples. Data from more than two groups were analyzed using multiple pairwise comparisons of the means through a one-way ANOVA including post-hoc tests corrected by a Bonferroni method. The significance threshold was set at 0.05 for all tests.

## Acknowledgments

The authors thank Lam Thuy Hoang, Marine Ibranosyan for help in clinical data management, Ken Bayles for help in providing Nebraska library mutants, Paul Verhoven for fruitful discussions, Laurent Schaeffer, Emilie Chopin and Nathalie Streichenberger from NeuroBio Tec-Banques (Hospices Civils de Lyon, France) for sharing expertise on muscle cells, CIQLE for technical assistance for microscopy, Katia Ziadna for all the agar plates and Laurence Cluzeau for excellent technical support. This work was supported by the EU grant INFECT-EU-FP7-HEALTH.

## Supplemental material

**Movie S1-S2. Murine myotubes.** Mature murine myotubes show spontaneous contractions.

**Fig. S1. Comparison of internalization of *Staphylococcus aureus* into murine myotubes and myoblasts.** The murine myoblast cells C2C12 were cultured in Dulbecco’s Modified Eagle’s Medium (Thermofisher) supplemented with 10% Fetal Bovine Serum (FBS) and 1% penicillin/streptomycin. Myoblasts were differentiated into myotubes by the addition of 2% horse serum in the medium for eight days. Internalization was determined after 2 hours of infection followed by antibiotic treatment to exclude extracellular bacteria. Percentage of numbers of viable bacteria was calculated in relation to infection inoculum. Multiplicity of infection (MOI) = 10. The diagram represents the mean ± standard deviations from three independent experiments of 6 NSTI-SA and 3 BSI-SA.

**Fig. S2. *Staphylococcus aureus* invasion is mediated by the eukaryotic fibronectin receptor α5β1 integrin.** Cells preincubated with monoclonal function-blocking antibodies were challenged with *S. aureus* strain RN6390ΔspA (A. Tristan, Y. Benito, R. Montserret, S. Boisset, E. Dusserre, F. Penin, F. Ruggiero, J. Etienne, H. Lortat-Jacob, G. Lina, M. Gabriela Bowden and F. Vandenesch, PLos ONE 4(4): e5042. doi:10.1371/journal.pone.0005042). This strain was chosen to avoid possible interference of Fc binding to staphylococcal protein (SpA) present in all *S. aureus* strains. Internalization was determined after 2 hours of infection followed by antibiotic treatment to exclude extracellular bacteria. Percentage of numbers of viable bacteria was calculated in relation to infection inoculum. Multiplicity of infection (MOI) = 10.

**Fig. S3. Expression of surface receptors in human keratinocytes and myoblasts**. Relative transcript levels were determined using quantitative reverse-transcriptase PCR, normalized to the internal *β-actin* standard and relative expression to human myoblasts. All values are means ± standard deviations of four independent experiments. The statistical significance was determined by Student’s t test with Welch’s correction (*p<0.05).

**Fig. S4. Cytotoxicity of staphylococcal or streptococcal culture supernatants on murine myoblasts or human keratinocytes.** Bacterial (NSTI-SA, 6 strains, blue lines; BSI-SA, 3 strains, brown lines; NSTI-GAS, 4 strains, green lines) supernatant (1:50 dilution) was added to the cell culture medium containing propidium iodide (PI) and cell death was quantified by monitoring PI incorporation over a 3 hours period. Triton X100 (red line) was used as positive control. Sterile bacterial culture medium (grey line) and cell culture medium (black line) were used as negative controls.

**Fig. S5. Phylogenetic analysis of the *S. aureus* strains.** Rooted phylogenetic analysis using a maximum likelihood approximation based on the 86233 SNPs identified through the conserved core genome of the 30 *S. aureus* strains. The tree is rooted on the clonal complex 1 (MSSA476). NSTI-SA isolates are labeled in red and BSI-SA in black.

**Text S1.** Detailed genomic analysis of the 30 *S. aureus* strains.

